# Molecular approaches reveal speciation between red and blue flowered plants in the Mediterranean *Lysimachia arvensis* and *Lysimachia monelli* (Primulaceae)

**DOI:** 10.1101/2021.04.16.440231

**Authors:** F.J. Jiménez-López, J. Viruel, M. Arista, P.L. Ortiz, M. Talavera

## Abstract

Flower colour constitutes a pivotal evolutionary force in speciation. The Mediterranean *Lysimachia arvensis* and *L. monelli* are morphologically variable species having both blue or red flowered plants. Previous studies suggested that *L. arvensis* plants differing in colour are diverging lineages, but this variation has not been considered in a phylogenetic context. We reconstruct the phylogenetic signal and the ancestral states of flower colour of Mediterranean *Lysimachia* species by using nuclear (ITS) and three plastid markers. Blue and red specimens are nested in two independent clades in the ITS tree, thus supporting that *L. arvensis* and *L. monelli* are polyphyletic, whereas low phylogenetic resolution was found in plastid markers. Blue-flowered *L. arvensis* is reconstructed sister to *L. talaverae* in a monophyletic clade sister to the remaining *Lysimachia*. Red-flowered *L. arvensis* is reconstructed sister to red-flowered *L. monelli* in a monophyletic clade sister to blue-flowered *L. monelli* and *L. foemina*. Our results suggest that colour lineages in *L. arvensis* and *L. monelli* constitute different species, but flower colour did not promote the separation of these lineages. We propose a new name for blue-flowered *L. arvensis* (*L. loeflingii*) and a new combination for red-flowered *L. monelli* (*L. collina*).

## Introduction

The systematic circumscription of the taxa belonging to the *Lysimachia* complex has been thoroughly studied since Linnaeus descriptions (Harborne, 1968; Källersjö, Bergqvist & Anderberg, 2000; Martins, Oberprieler & Hellwig, 2003; Hao et al., 2004; Manns & Anderberg, 2007a, 2007b; Zhang et al., 2012). Traditionally, *Anagallis* L. had always been considered a genus closely related to *Lysimachia* L. However, phylogenetic analyses based on the nuclear ITS and plastid data resolved species belonging to either *Anagallis* and *Lysimachia* intermingled across different clades (Manns & Anderberg, 2005). For example, the Mediterranean *A. arvensis* L. and *A. monelli* L. were more closely related to species of *Lysimachia* and other genera such as *Pelletiera* A.St.-Hil., *Asterolinon* Hoffmanns. & Link, and *Trientalis* L., than to other *Anagallis* species (Anderberg, Manns, & Källersjö, 2007). These phylogenetic results led to the inclusion of some taxa previously included in *Anagallis* within *Lysimachia* (Manns & Anderberg, 2007b, 2009); and new combinations were subsequently proposed for *L. arvensis* (L.) U. Manns & Anderb. and *L. monelli* (L.) U. Manns & Anderb. (Manns & Anderberg, 2009).

All these systematic readjustments are congruent with the high phenotypic variability found within the former genus *Anagallis*, in which morphological traits, such as leaf shape or colour, size and presence of glands in flowers, hinder the definition of taxonomic boundaries. The Mediterranean *L. arvensis* and *L. monelli* show a high morphological and ecological diversity, and consequently a wide number of infraspecific taxa have been described within each species. For example, diploid plants (2*n*=20; Šveřepová, 1972; Kress, 1969; García Pérez et al., 1997) with small size and small flowers, and specific ecology (Bolòs & Vigo, 1996; Gibbs & Talavera, 2001) originally recognized as *A. parviflora* Hoffmanns. & Link (Hoffmannsegg & Link, 1813-1820), were classified as infraspecific taxa of *L. arvensis*, or currently recognized as *L. talaverae* L. Saéz & Aymerich (Aymerich & Sáez, 2015). Likewise, plants morphologically very similar to *L. arvensis* but with four cells in the marginal glands of the petals (instead of three) were first recognized as *A. arvensis* subsp. *foemina* (Mill.) Schinz & Thell., and later described as a different species: *L. foemina* (Mill.) U. Manns & Anderb. (Manns & Anderberg, 2009). This taxonomic reorganization was based on phylogenetic results that reconstructed *L. monelli* as sister to *L. foemina* (Manns & Anderberg, 2007a). Several taxa have also been recognized from the infraspecific variability within *L. monelli*, such as plants with narrow and linear leaves which were considered as *L. monelli* subsp. *linifolia* (L.) Peruzzi (Peruzzi, 2010). Likewise, plants with small and wide leaves and short internodes were recognized as *L. monelli* subsp. *maritima* (Mariz) Carlón et al. (Carlón et al., 2014).

Both *L. arvensis* and *L. monelli* show flower colour polymorphism with blue and red-flowered plants (Ferguson, 1972; Pujadas, 1997). This trait further increased the taxonomic complexity because numerous taxa have been described in relation to flower colour. The original description of *A. arvensis* was based on a red-flowered specimen (Linnaeus 1753), whereas the blue morph was first described as *A. latifolia* L. (Linnaeus 1753), then as *A. caerulea* L. (Linnaeus 1759) but it is now accepted as *L. arvensis* var. *caerulea* (L.) Turland & Bergmeier (Bergmeier et al., 2011). Blue and red plants of *L. arvensis* show different geographical distributional pattern, pure blue or mixed populations appear chiefly in dryer Mediterranean localities while pure red populations predominate in more temperate areas (Arista et al., 2013). In addition, studies using microsatellite markers showed the separation of both colour morphs in two groups (Jiménez-López et al., 2019a). Both colour lineages of *L. arvensis* differ in other traits as flower phenology or type of herkogamy (Arista et al., 2013; Jiménez-López et al., 2020), and salmon-flowered plants resulting from the cross between red and blue individuals are rare in mixed populations (Jiménez-López et al., 2020).

In the case of *L. monelli*, the species description was based on a blue-flowered specimen (Linnaeus 1753), and the red-flowered individuals were first described as *A. collina* Schousb. (Schousboe, 1800), then recognized by various authors as infraspecific taxa of *Anagallis monelli* or *A. linifolia* L. (Linnaeus, 1762) (see taxonomic implications). Currently, these names are accepted within *Lysimachia monelli*. Populations of *L. monelli* with blue-flowered plants are restricted to the Iberian Peninsula, Morocco, Algeria, northwest of Libia (Tripolitania) and Tunisia (Jahandiez & Maire, 1934) while the red-flowered plants grow mainly in Morocco, Sardinia and northeast of Spain (Willkomm, 1870; Jahandiez & Maire, 1934).

The huge morphological variability of the Mediterranean *Lysimachia* complex is currently reduced to four species: *L. arvensis, L. monelli, L. foemina* and *L. talaverae* (Manns & Anderberg, 2009; Aymerich & Sáez, 2015). *Lysimachia arvensis* is a Mediterranean species that at present is distributed across the world. It is annual, self-compatible (Gibbs & Talavera, 2001) and tetraploid (2*n*=40; revised in Pastor, 1992; García Pérez et al., 1997; Monein, Atta & Shehata, 2003). *Lysimachia monelli* is distributed exclusively in the west of the Mediterranean Region. It is perennial, self-incompatible (Gibbs & Talavera, 2001; Talavera et al., 2001; Freyre & Griesbach, 2004) and diploid (2*n*=20; Kress, 1969; Valdés, 1970; Šveřepová, 1972; García Pérez et al., 1997). *Lysimachia foemina* is also a Mediterranean species but, as *L. arvensis*, at present is distributed across the world. It is annual, self-compatible (Marsden-Jones, 1935; Marsden-Jones & Weiss, 1938) and tetraploid (2*n*=40; revised in Pastor, 1992). *Lysimachia talaverae* is endemic to the western Mediterranean, it is annual, self-compatible (Gibbs & Talavera, 2001) and diploid (2*n*=20; Šveřepová, 1972; García Pérez et al., 1997).

Previous phylogenetic studies have scarcely explored the potential implications of colour polymorphism in taxa delimitation because it was considered as part of the intraspecific variation, and usually only one colour per species was sampled in molecular analyses (Martins et al., 2003; Manns & Anderberg, 2005; Anderberg et al., 2007; Manns & Anderberg, 2007b; Yan et al., 2018). However, Manns & Anderberg (2007a) reconstructed a phylogenetic tree including samples of both colours of *L. arvensis* from a single population, which were resolved as monophyletic using plastid markers, but appeared in independent clades in the ITS-based phylogenetic tree. All these evidences suggest that larger sampling of the two colour lineages for both *L. arvensis* and *L. monelli* is required in phylogenetic studies to clarify the implications of flower colour in taxa boundaries. In this context, our study aims to (1) define the phylogenetic identity of the blue and red lineages of *L. arvensis* and *L. monelli*, (2) establish the role of flower colour in the divergence of these taxa and (3) propose new names or combinations in accordance with the results of phylogenetic analyses where necessary.

## Material & methods

### Plant material

Freshly leaf material from 28 populations of *L. arvensis*, 12 of *L. monelli*, three of *L. foemina* (LF) and four of *L. talaverae* (LT) (Fig. 1, Table S1) were collected and dried in silica-gel (Chase & Hills, 1991). This sampling represents nine pure red and 12 pure blue populations and seven mixed (red and blue) populations of *L. arvensis* (LA_R and LA_B, respectively). Sampling of *L. monelli* included four pure red (LM_R) and eight pure blue populations (LM_B). Also, one sample of *L. azorica* Hornem. ex Hook. (LZ), and one sample of two populations of *L. linum*-*stellatum* (L) Hoffmanns. & Link (LLS). Two additional related species were included in the phylogenetic analyses and used as outgroups (Table S1).

**Figure 1.**
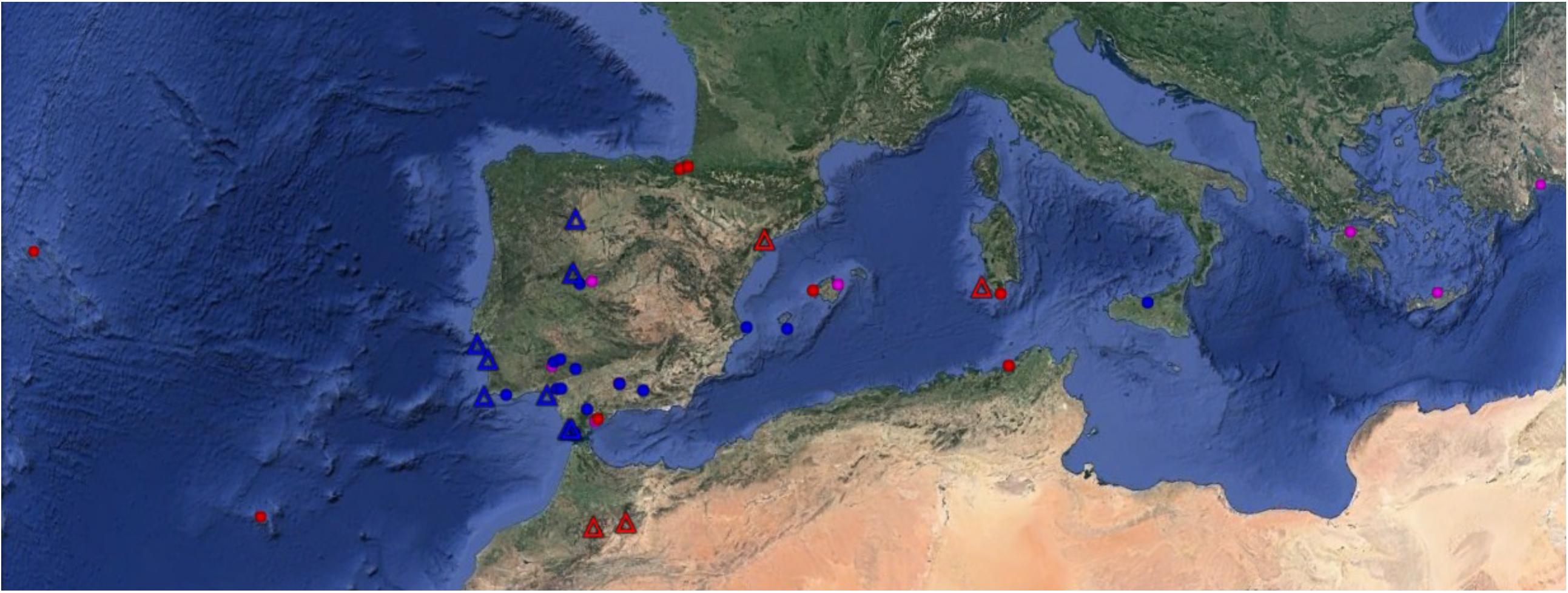
– Geographical distribution of the studied populations of *Lysimachia arvensis* (circles) and *Lysimachia monelli* (triangles). Colours correspond to flower colour of the studied populations: blue and red for pure blue and red populations, respectively, and pink for mixed populations.

### DNA isolation, amplification and sequencing

Total genomic DNA was isolated with Invisorb Vegetal DNA Kit HTS 96 (Invitek, Spain), with modifications following Jiménez-López et al. (2016). The quality of the extracted DNAs was checked in 1% TAE-agarose gel, and DNA concentration was estimated using a NanoDrop DS-11 Spectrophotometer (DeNovix).

Both ITS and plastid markers were used for phylogenetic reconstructions. The ribosomal ITS region partial18S–5.8S–partial28S was amplified using ITS5 and ITS4 primers published by White et al. (1990). We tested 19 plastid markers in eight samples of *L. arvensis* (four blue and four red specimens), but only three of them were polymorphic. The polymorphic plastid markers *rps16*–*trnK, rpl32-trnL* were amplified following Shaw et al. (2007), and *trnH*–*psbA* following Hamilton (1999). PCR amplifications were performed following Jiménez-López et al. (2016) using 4ng of DNA as input and the amplified fragments were checked in 2% TAE-agarose gels. An Applied Biosystems VeritiTM Thermal Cycler was used with the following programmes: for *rps*16-*trn*K and *rpl*32*-trn*L: first step, one cycle of 80 °C for 5 min; a second step, 35 cycles with three consecutive conditions: 95 °C for 1 min, increase from 50 to 65 °C during 1 min and 65°C for 3 min; third step, one cycle of 65°C for 5 min and then 4 °C. For *trn*H-*psb*A the conditions were first step, one cycle of 96 °C for 5 min; second step, 35 cycles with three consecutive conditions: 95 °C for 45 sec, 53 °C for 1 min and 72 °C for 30 sec; third step, 1 cycle of 72 °C for 5 min and then 4 °C. After purification with ExoSAP-IT (Roche, Spain), fragments were sequenced in an ABI 3730 machine, BigDye Terminator Cycle Sequencing Kit (Applied Biosystems, Foster City, CA, USA), at STAB Vida Lda. (Oeiras, Portugal). Sequences were edited using Geneious R10 (Biomatters Ltd, Auckland, New Zealand). Alignments were conducted in MEGA ver. 6 (Tamura et al., 2013) using the algorithm Clustal IW (Thompson, Gibson & Higgins, 2002), and manually edited.

### Phylogenetic inference

The Akaike information criterion (AIC) was used to estimate the evolution model that better fit each DNA matrix using jModeltest 2.1.4 (Darriba et al., 2012). Phylogenetic relationships were inferred based on maximum parsimony inference using PAUP ver. 4.0b 10 (Swofford, 2003) and maximum likelihood methods using RAxML Ver. 8 (Stamatakis, 2014). Bootstrap support (BS) was estimated with 500,000 bootstrap replicates following full heuristic searches. Bayesian inference was conducted using MrBayes on X_SEDE_ (3.2.7a) (CIPRES Science Gateway; Miller, Pfeiffer & Schwartz, 2010) and BEAST v.2.5 (Bouckaert et al., 2019). We use coalescent models (Heled & Drummond, 2010) and we ran 30 independent analyses with 50 million generations, sampling every 5000. The phylogenetic reconstructions were calculated for each plastid and nuclear markers independently, and for a combined dataset including the three plastid datasets. Following Viruel et al. (2016) clades with bootstrap support (BS) of 75-100 or posterior probabilities (PPS) of 0.95-1.0 were considered moderately to strongly supported. Phylogenetic reconstruction with ITS was performed with and without the sample LA19_B according to the results of the recombination analysis (see below and results section).

In order to test for incongruent topologies between the independent DNA data matrices, a partition homogeneity test was performed (Farris et al., 1994, Symonds & Lloyd, 2003). The Incongruence Length Difference (ILD) test of Farris et al. (1994) implemented in PAUP ver.4.0b 10 (Swofford, 2003) was performed through 1,000 random-order-entry replicates to estimate if the nuclear and the three plastid data sets were significantly different from random partitions of the same size.

### Recombination Analysis

To test possible recombination events in the ITS sequences of all the samples, the seven methods (RDP, GENECONV, BootScan, MaxChi, Chimaera, SiScan & 3SEQ) implemented in RDP4 v.484 were applied to determinate parental and recombinant sequences with their probability scores for hypothetical recombination events (Martin et al., 2010; Viruel et al., 2018).

### Ancestral state reconstruction

To estimate the basal state of flower colour, we performed ancestral state reconstruction of colour classes using stochastic character mapping (SIMMAP; Bollback, 2006) with the package ‘phytools’ (Revell, 2012) implemented in R (R Core Development Team, 2015). We performed the analyses on the RAxML phylogenetic tree obtained from ITS dataset. Because we included more than one sample per species, we also pruned the additional samples for the same species in the case of monophyly. For SIMMAP analyses, we ran 1000 simulations per tree. Ancestral state reconstructions for all additional nodes were estimated with parsimony and SIMMAP methods only. Ancestral states reconstruction was shown only on major clades.

### Divergence-time estimation

In addition, we used molecular clock approaches to estimate divergence times among the main clades. We selected one representative sequence per species, together with sequences of other *Lysimachia* and related species from GenBank (Table S1).

In this case, the phylogenetic reconstruction was inferred with maximum likelihood methods with RAxML (Stamatakis, 2014). We used a Bayesian uncorrelated-lognormal relaxed-clock approach (Drummond et al., 2006) as implemented in BEAST v.2.5 (Bouckaert et al., 2019). Node ages and absolute branch substitution rates were estimated using two relaxed clock methods that represent two contrasting approaches to estimate divergence times (Magallón, Hilu & Quandt, 2013). We used both Yules speciation and birth–death tree priors (Heled & Drummond, 2012). We ran 30 independent analyses (50 million generations each, sampled every 5000). Tracer 1.7.1 software (Rambaut et al., 2018) was used for convergence and satisfactory Effective sample size (ESS) values (>200) and verify that a burn limit of 10% was appropriate. Subsequent parameter distributions were obtained by combining the independent MCMCs with LogCombiner 1.10.4, and calculating the maximum credibility tree using TreeAnnotator 1.10.4 (Rambaut et al., 2018).

Fossil records are scarce because these plants lack woody parts, and both seed and pollen production are low. The genus *Maesa* Forssk is at the root of the *Primulaceae* phylogeny (Anderberg et al., 2007; Yesson, Toomey & Culham, 2009) and its pollen is known from the Early Eocene (55.8 Ma; Brook, Burney & Coward, 1990; Kershaw, Bretherton & Kaars, 2007) according to The Paleobiology Database (http://paleodb.org/). Furthermore, fossil seeds similar to those of *Lysimachia vulgaris* L. and its close relatives (Oh et al., 2008) are found from the Middle Miocene (12–16 Ma; Friis, 1985). We therefore used the pollen record to set the age prior for rooting and seed fossil to set the age prior for *L. vulgaris*. A secondary calibration was applied to the crown age of core *Lysimachia* using an estimated mean age of 30 Ma from previous phylogentics reconstructions (Yesson et al., 2009, Zanne et al., 2014). The most generally appropriate prior for fossil calibrations is the lognormal distribution (Ho & Phillips, 2009). Then, for the crown age of core Lysimachia node we used a standard lognormal prior distribution with mean of 30 Ma and standard deviation (SD) of 4 Ma following Yang et al. 2018. In order to test the effect of the different calibration points and tree prior approaches, we also performed analyses using only one fossil calibration, only secondary calibration and both calibration points. We then performed a regression between the node ages of same clades from all analyses.

## Results

### Phylogenetic reconstructions and chronogram inference

Fifty-two high quality sequences were obtained for ITS, and 37 for the plastid regions. The pDNA regions were 675-750 bp long for *rps16*–*trnK*, 580-630 bp for *rpl32*-*trnL* and 365-445 bp for trnH-psbA. The ITS1 and ITS2 fragments ranged from 600 to 737 bp. The G+C content in all samples was 22.22% for *rps16*–*trnK*, 27.25% for *rpl32*–*trnL*, 29.04% for *trnH-psbA* and 57.88% for ITS on average. The G+C contents were significantly different between *L. arvensis* and *L. monelli* for ITS (*p*=0.001; 58.86% and 56.46%, respectively) and for rps16-trnK (*p*=0.015; 22.17% and 22.30%, respectively). No significant differences were found on the G+C contents among blue and red individuals of *L. arvensis* (*p*=0.482) or *L. monelli* (*p*=0.338).

The parsimony, RAxML and Bayesian trees reconstructed the same topology for all analysis for the ITS region (Fig. 2). Likewise, for each pDNA region separately and for the combined matrix, the tree reconstructed showed a very similar topology in all analyses and markers (Figs. 3). However, the ILD test showed significant differences between nuclear and plastid regions (*p*=0.010), indicating that there was an inconsistency between these markers. Thus, phylogenetic analyses were performed for the ITS region separately from the concatenated dataset representing the three plastid markers.

**Figure 2.**
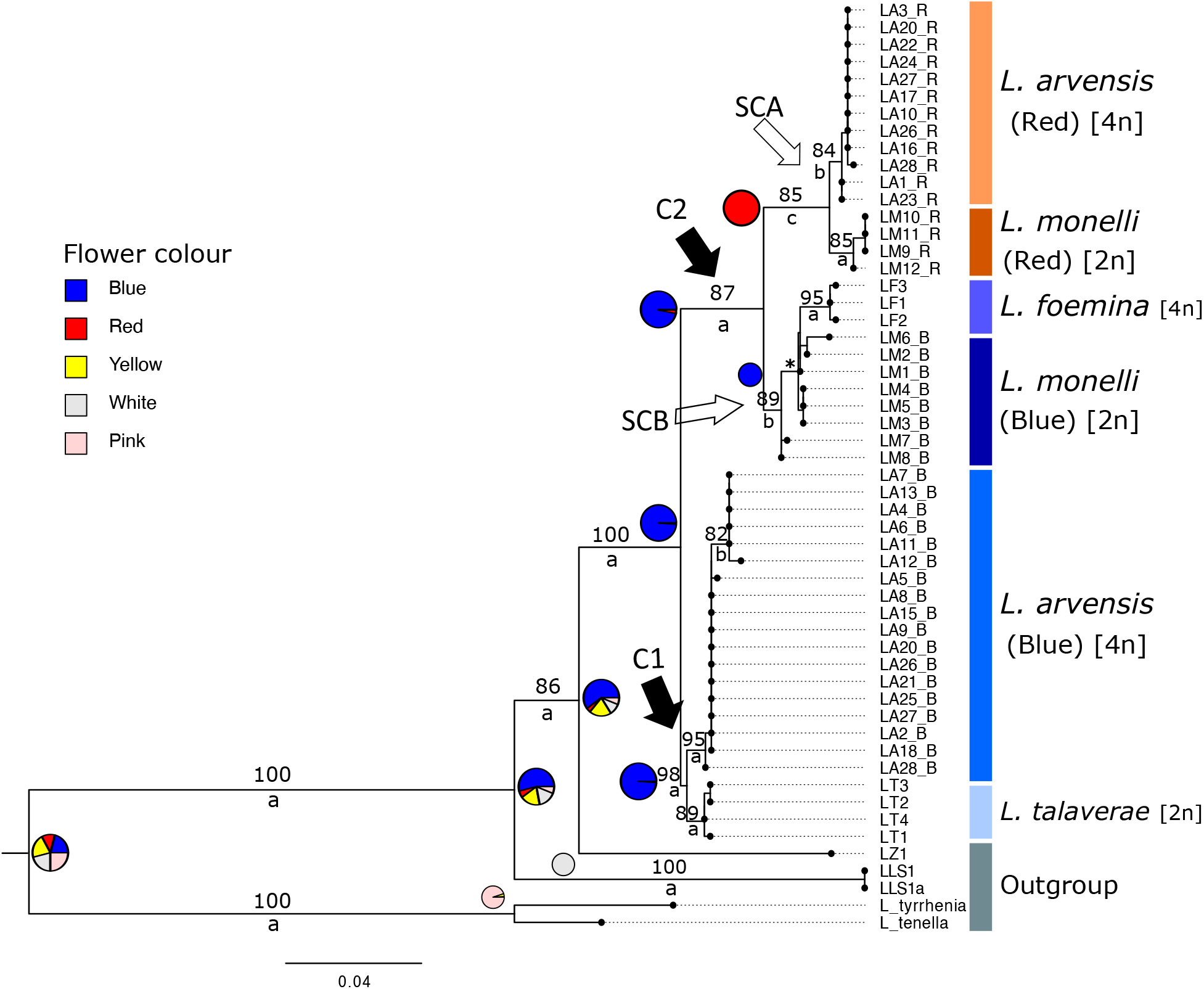
– Ancestral state reconstruction of flower colour and ploidy level of *L. arvensis* and *L. monelli* and related taxa (see Table S1). RAxML tree based on ITS sequences. The samples of *L. arvensis* from mixed populations are highlighted in bold. Bootstrap values > 50% are given above the branches and Posterior probabilities of Bayesian approach are given below branches: a (0.995-1.000 PPS), b (0.950-0.994 PPS) and c (0.850-0.949). C1: Clade1; C2: Clade 2; SCA: Subclade A and SCB: Subclade B. * (83 BS, 0.984 PPS).

**Figure 3.**
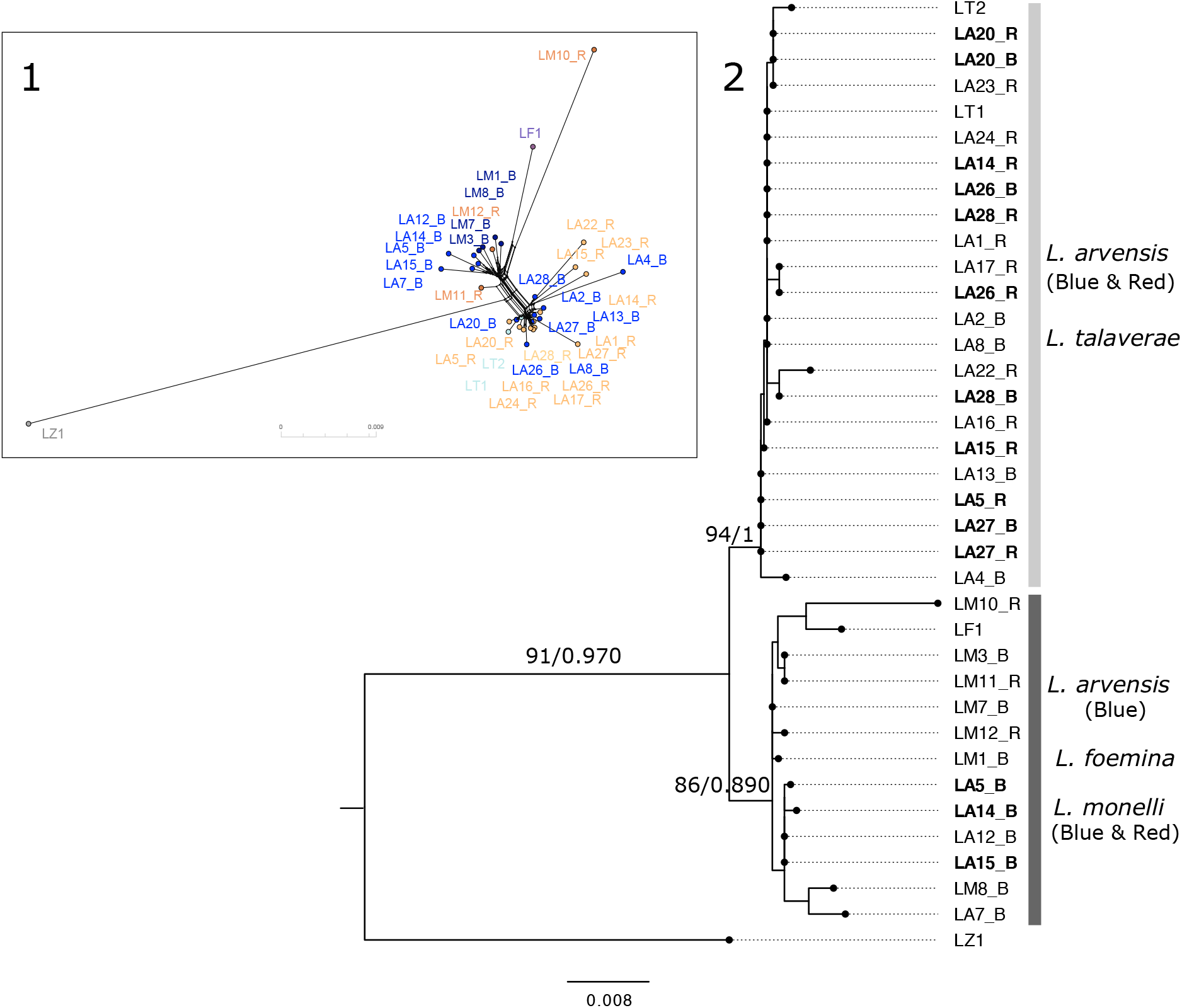
– (1) Phylogenetic network and (2) Maximum likelihood (RAxML) tree based on concatenated plastid sequence data (*trn*H–*psb*A, *rps*16– *trn*K and *rpl*32-*trn*L) of *L. arvensis, L. monelli* and related taxa (see Table S1). Main clades in trees from ML (RAXML) and Bayesian analysis (MrBayes) are identical. Bootstrap values > 50% and posterior probabilities of Bayesian tree are given above and below the branches respectively. Samples of *L. arvensis* from mixed populations are highlighted in bold.

The phylogenetic tree reconstructed with the concatened pDNA matrix showed two clades (Figs. 3). The first clade (94 BS, 1.000 PPS) comprised all the red and some blue individuals of *L. arvensis* and *L. talaverae*. The second clade (86 BS, 0.890 PPS) included blue individuals of *L. arvensis, L. monelli* (blue and red) and *L. foemina*. This topology was also observed for each plastid marker independently, although the support bootstrap was lower in all cases.

The RAxML tree constructed with ITS showed two main clades (Fig. 2). In the Clade I (C1, 98 BS, 1.000 PPS), *L. arvensis* blue samples (95 BS, 1.000 PPS) were sister to *L. talaverae* (89 BS, 0.999 PPS), whereas the samples of *L. arvensis* from SW of Spain formed a clade (82 BS, 0980 PPS). The Clade II (C2, 87 BS, 1.000 PPS) comprised two subclades, named A and B. In the subclade A (SCA, 85 BS, 0.935), red *L. arvensis* samples (84 BS, 0.976 PPS) were sister to the red *L. monelli* samples (85 BS, 1.000 PPS). In the subclade B (89 BS, 0.990), blue *L. monelli* samples (83 BS, 0.984 PPS) were sister to *L. foemina* (95 BS, 1.000 PPS). Therefore, blue and red individuals of both *L. arvensis* and *L. monelli* appeared in different groups. It is noteworthy that blue and red individuals from mixed populations of *L. arvensis* appeared in independent clades (Figs. 2). In addition, most of the ancestral reconstruction states showed high probability of blue-flowered plants until the SCA, where red flowers appeared (Fig. 2).

Similar time estimates and topologies were obtained using the three different calibration ages and different tree priors for the stem age of *Lysimachieae* tribe (Fig. 4). Divergence time results based on these different strategies were all very similar to each other. Our primary analyses (Fig. 4) used the full tree with the Yule prior and all calibration points. The correlation between the node ages of clades from the primary tree and alternative trees based on different calibration schemes and tree priors were significant for all cases (r^2^ = 0.978-0.891, p=0.000) The maximum clade credibility (MCC) tree of *Lysimachiae* tribe showed three clades in *Lysimachia* (Fig. 4): one with *L. arvensis, L. monelli* and related species (BS 100%; 1.000 PPS, Clade A), other with *L. vulgaris, L. punctata* and related species (0.940 PPS, Clade B), and a third clade of *Lysimachia*, with *L. tenella* or *L. minima* and other related species (BS 100%; 0.990 PPS, Clade C). The latter clade appeared closest to *Cyclamen* sp. (BS 100%; 1.000 PPS). According to the relaxed molecular clock and Yules tree priors, the common ancestor of blue plants of *Lysimachia arvensis* and *L. talaverae* diverged ca. 1.5 Mya (95% HPDs 0.2-2.5), and the divergence of blue plants of *L. monelli* and *L. foemina* likely occurred ca. 0.6 Mya (95% HPDs 0.1-1.6) and the split between red plants of *L. arvensis* and red plants of *L. monelli* was estimated ca. 1.2 Mya. (95% HPDs 0.2-2.4). Furthermore, the common ancestor between *L. arvensis* red-*L. monelli* red and *L. monelli blue/L. foemina* subclades likely diverged ca. 1.4 Mya (95% HPDs 0.6-2.9) and the divergence between these clades and *L. arvensis*/*L. talaverae* clade occurred ca. 3.6 Mya (95% HPDs 1.3-7.5) (Figs. 4 and S7). Two events of divergence occurred during Miocene period between the different clades of *Lysimachia*. The first one happened about 28.9 Mya (95% HPDs 16.9-47.2), when diversifying the clade C from the other *Lysimachia* clades, and the second one happened about 24.2 Mya (95% HPDs 13.1-42.6) dividing clades A and B (Fig. 4).

**Figure 4.**
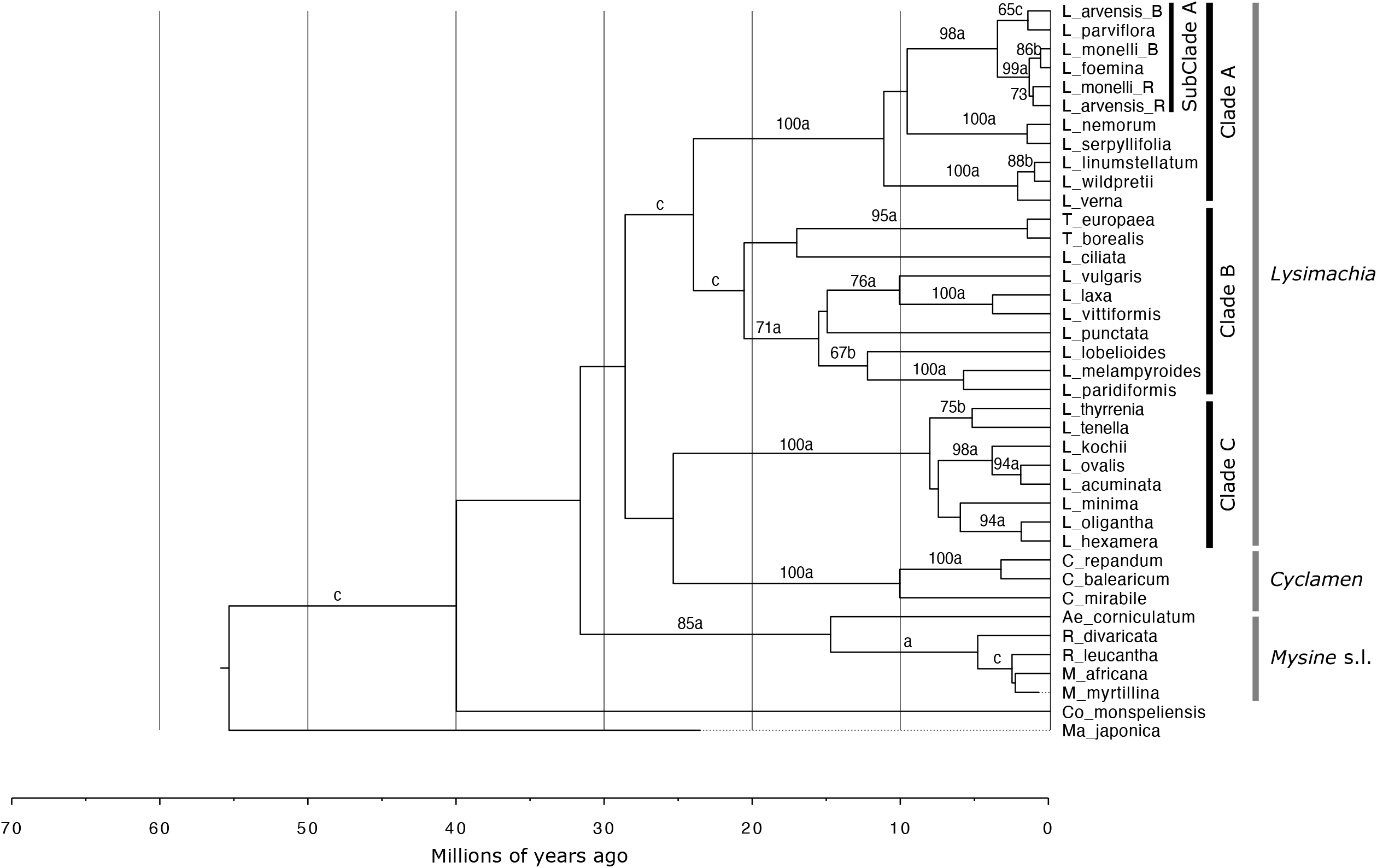
– Phylogenetic reconstruction for Lysimachiae tribe. Maximum Clade Credibility (MCC) tree reconstructed with BEAST based on Internal transcribed spacer (ITS) data of *Lysimachia* species and related genera of tribe Lysimachieae. The root was calibrated following Yan et al. 2018 and a *Maesa* sp. fossil (Kershaw et al. 2007), and a Yules speciation tree prior. Bootstrap support > 65% of ML estimate (RAxML) and Posterior probabilities of Bayesian approach (BEAST; a [0.995-1.000 PPS], b [0.950-0.994 PPS] and c [0.850-0.950]) are given above branches.

### Recombination Analyses

Four of the seven recombination methods detected two recombinant regions in the sample LA19_B (*L. arvensis* blue). An insertion of 18 nucleotides at position 16 from *L. arvensis* and an insertion of 25 nucleotides at position 133 from *L. foemina* were observed. Therefore, this sample was discarded from the tree in Figure 2.

## Discussion

Our phylogenetic results based on the nuclear ITS region demonstrates that blue and red individuals of *Lysimachia arvensis* and *L. monelli* are independent taxa (Fig. 2). The presence of different ITS sequences in the red and blue flowered individuals of *L. arvensis* collected from the same populations had already been reported by Manns & Anderberg (2007a). The high support of our ITS results (89 BS, 0.999 PPS) suggests that blue and red lineages of *L. arvensis* belong to different taxa, and in addition, both colour taxa are much more close to other species that between them. On one hand, the blue flowered-plants of *L. arvensis* are sister to *Lysimachia talaverae* (Clade I). This last species has already been recognized with the rank of species based only on morphological, ecological and kariological traits (Aymerich & Sáez, 2015) but not on molecular tools. Our study further validates the species rank status of *L. talaverae* using phylogenetic inference. On the other hand, red individuals of *L. arvensis* are sister to red individuals of *L. monelli* (Subclade A). Likewise, the two colour lineages of *L. monelli* should be also considered as different taxa, being the blue lineage of *L. monelli* sister to *L. foemina* (Subclade B), supporting the independence of *L. foemina* from *L. arvensis*, in accordance with Manns & Anderberg (2007a).

Colour lineages of both species appeared together with pDNA markers. Hybridization has been proposed as causes of incongruences between pDNA and nuclear phylogenies (Wendell & Doyle, 1998;> Semerikova & Semerikov 2016). Hybridization involves past or present contact or hybrid zones (Petit, Bretagnolle & Felber, 1999), where lineages with divergent genomes have the potential to exchange genes (Souissi et al., 2017). However, as a result of this exchange it would be difficult to find a clear separation between lineages, contrasting with the strong separation between *Lysimachia* species using the nuclear ITS (Fig. 2). Only one sample (LA19B, blue *L. arvensis*) showed hybridization evidence according to the recombination analyses. The position of this sample in the ITS tree would suggest a possible hybridization between blue *L. arvensis* and *L. foemina*. These two taxa co-occur in some populations in Europe and their flowers are very similar, thus they possibly share pollinators. However, the recombination analysis pointed that this hybrid resulted from the crossing between red *L. arvensis* and *L. foemina*. Thus, that hybridization was originated from an individual with blue flowers very similar to the blue *L. arvensis* (as it was initially considered in the sampling). This hybrid (*L. arvensis* x *L. foemina*) has already been described as *A*. x *doerfleri* Ronniger (Dörfler, 1903). (=*Lysimachia* x *doerfleri* (Ronninger) Stace). However, hybrid natural populations stabilisation seems difficult, since experimental crosses carried out by other authors frequently resulted in sterile F_1_ progeny (Marsden-Jones, 1935; Kollmann & Feinbrun, 1968; Šveřepová, 1972).

Our results corroborate the maternally haplotype inheritance of the plastomes (Wolfe, Li & Sharp, 1987), and their sequences are frequently identical among closely related species in many plant taxa (Wolf, Murray & Sipes, 1997; Matsumura et al., 2009). The incongruent pattern observed between the two phylogenetic trees (Degnan & Rosenberg, 2009) can be also explained by the coalescent theory (Nordborg, 2001; Wakeley, 2009) because two or more lineages can coexist in the same ancestral population (Degnan & Rosenberg, 2009). Thus, the common ancestor of this group would have had the two pDNA haplotypes, which are currently present in one of them in red individuals of *L. arvensis* and *L. talaverae*, the other in *L. monelli* and *L. foemina*, and both haplotypes in blue individuals of *L. arvensis*. Later, a segregation of colour lineages of *L. arvensis* and *L. monelli* likely happened. That segregation could have been promoted by geographic separation, assortative mating mediated by pollinators or by differential tolerance to abiotic factors as has been described in other species (Kirkpatrick, 2000; Strauss & Whittal, 2006; Hopkins & Rausher, 2012; Wang et al., 2013). In *L. monelli* geographic separation could be contributing to lineage separation. However, in *L. arvensis* genetic flow is potentially possible between lineages in mixed populations although the reproductive isolation between them is high (Jiménez-López et al., unpublished data); in fact, the results of nuclear phylogenetic reconstruction points towards a clear isolation of both colour lineages.

Flower colour constitutes a pivotal evolutionary force to speciation in several groups of plants (Carlson & Holsinger, 2015; Ellis & Field, 2016; Takahashi, Takakura & Kawata, 2016, Narbona et al., 2018) and it has been proposed as a “magic trait”, that is a trait “encoded by genes subjected to divergent selection that affect pleiotropically reproductive isolation” (Servedio et al., 2011). However, according to ancestral state reconstruction, flower colour does not seem the trait promoting divergence between lineages of *L. arvensis* or in those of *L. monelli*. Ancestral states reconstruction of flower colour points to a common blue-flowered ancestor of this group and the transition to red-flowered plants occurred only once, at the divergence node of red *L. arvensis* and *L. monelli*. This kind of transition from blue to red in flowers is quite frequent due to the inactivation of a branch of the anthocyanin pathway (Rausher, 2008). In other plant groups, red-flowered species usually derived from blue-flowered species (Kay et al., 2005; Wilson et al., 2007; Rausher, 2008; Wessinger & Rausher, 2012). In our study, the blue ancestor likely originated two lineages, one entirely blue (which includes the blue lineage of *L. arvensis*) and another that subsequently separated into a blue and another red subclade (which includes the red lineage of *L. arvensis* and *L. monelli*). This result indicates that flower colour cannot have been the trigger for the current speciation of *L. arvensis* or *L. monelli* colour lineages.

The location of each colour lineage of *L. arvensis* and *L*.*monelli* in the phylogenetic reconstruction suggests independent origins for colour lineages of both species. According to the phylogenetic tree constructed with ITS, both a diploid and a tetraploid taxa appear together in each clade: blue lineage of *L. monelli* (2*n*) with *L. foemina* (4*n*), *L. talaverae* (2*n*) with blue lineage of *L. arvensis* (4*n*), and red lineage of *L. monelli* (2*n*) with red lineage of *L. arvensis* (4*n*). That configuration strongly suggests the role of polyploidy in the speciation process of this complex. Frequently speciation events in flowering plants involve hybridization processes and subsequent modification of divergent genomes (Stebbins, 1971). Polyploid speciation implies the complete duplication of the genome after hybridization (allopolyploidy). In addition, new species can be produced not by hybridization, but by duplication of complete genomes which also leads to polyploidy (autopolyploidy). The configuration observed in the phylogenetic reconstruction suggests that both colour lineages of *L. arvensis*, which are tetraploid, have been originated by autopolyploidy. Red *Lysimachia arvensis* could derive from red *L. monelli* (diploid) and blue *L. arvensis* could derive from *L. talaverae* (diploid with blue flowers). Moreover, *L. foemina* (tetraploid with blue flowers) could derive from blue *L. monelli* (diploid). Although this is a hypothesis, at least for *L. arvensis*, both colour lineages present a karyotype with four equal copies from a set of chromosomes (Monein et al., 2003), supporting the autopolyploid origin of these taxa. This would indicate the presence of multiple polyploidization events in the clade studied, something that has previously been found in other genera (eg. Rieseberg et al., 2003; Wang et al., 2006; Wood et al., 2009; Soltis et al., 2015; Padilla-García et al., 2018). Divergence-time estimation between the diploid and tetraploid taxa, ca 1Mya (Fig. 4) and the support of the phylogenetic reconstruction (Fig. 2) point to non-recent tetraploidization events, and the stability in the genomic sequences studied also suggest an autopolyploid origin, in accordance with current evolutionary theories on polyploids (Otto & Whitton, 2000; Soltis, Visger & Soltis, 2014; Soltis et al., 2015). However, this role of polyploidy in the evolution of the studied *Lysimachia* would need a more directed study.

In conclusion, the two colour morphs of *L. arvensis* constitute independent lineages and their current isolation is evident. Blue *L. arvensis* is a more adapted taxon to the arid conditions of the Mediterranean while red lineage is a more mesic taxon with a greater distribution area, co-occurring with the blue lineage in some Mediterranean areas (Arista et al., 2013). These two colour lineages also separate when using nuclear microsatellite markers (Jiménez-López et al., 2019a). Similarly, the two colour lineages of *L. monelli* are also independent entities with allopatric distribution. A phylogeographic approach would be necessary to clarify the geographical origin of both colour lineages. Although our results indicated that flower colour has not triggered recent speciation in this group, it can have an important role as a reinforcement mechanism that prevents the formation of maladapted hybrids (Hopkins, 2012). This could be the situation at least in *L. arvensis*, where pollinators prefer the blue lineage and barely made transitions between flowers of different colours (Ortiz et al., 2015; Jiménez-López et al., 2019b). The low frequency of hybrids in mixed-colour populations (Jiménez-López et al., 2020) and the lack of hybrid samples found in the present study support the importance of colour as reinforcement mechanism.

### Taxonomic implications

Our results have taxonomic implications for the colour lineages of *L. arvensis* and *L. monelli* as each lineage should be defined as different taxa with morphological, genetic and geographic identity. Thus, we propose the following names or combinations in accordance with molecular analyses:

-Red plants of *Lysimachia arvensis* should maintain this name because Linnaeus in 1753 described the species from red-flowered plants.

*Lysimachia arvensis* (L.) U. Manns & Anderb. in Willdenowia, 39(1):51. (2009) ≡ *Anagallis arvensis* L., Sp. Pl.: 148. 1753, [basion.] ≡ *Anagallis arvensis* var. *phoenicea* Gouan, Fl. Mons.: 24 (1764), nom. Illeg. ≡ *Anagallis phoenicea* (Gouan) Scop., Fl. Carn. ed. 2, 1: 139 (1777), nom. Illeg. ≡ *Anagallis arvensis* subsp. *phoenicea* (Gouan) Wallmann in Ber. Bayer. Bot Ges. 9: 44 (1904), nom. inv. Ind. Loc.: “Habitat in Europae arvis”.

Lectotype designated by Dyer & al. 1963: 14: Herb. Linn. 208.1 (LINN)

-Blue plants of *Lysimachia arvensis* should be called with the specific epithet latifolia as it was the first name employed by Linnaeus in 1753 for plants with blue flowers. However, the epithet latifolia already exists within *Lysimachia* referred to a different taxon [*Lysimachia latifolia* (Hook.) Cholewa in Phytoneuron 28: 1-2 (2014) = *Trientalis latifolia* Hook., Fl. Bor. Amer. 2(9): 121 (1839), a plant described from Washington]. Therefore, we have selected the name *Lysimachia loeflingii* because Linnaeus in 1753 described blue-flowered plants from materials collected by Loefling from Spain.

### Lysimachia loeflingii F.J. Jiménez-López & M. Talavera, nom. nov

*Anagallis latifolia* L., Sp. Pl. 1: 149 (1753) [syn. subst.] ≡ *A. arvensis* subsp. *latifolia* (L.) Arcang., Comp. Fl. Ital., ed.2: 456 (1894) - *Anagallis arvensis* L. subsp. *arvensis* var. *caerulea* sensu Kollmann & Feinbrun in Notes Royal Bot. Gard. Edingurg 27: 176 (1968), non *A. arvensis* L. var. *caerulea* (L.) Gouan, Fl. Monsp.: 30 (1765)

Ind. loc.: “Habitat in Hispania Loef.”

Lectotype designated here: Herb. Linn. n° 208.3, “H.U.3. latifolia. ex Hispanica foly amplexicauly” [m. Linnaeus]

*Anagallis latifolia* L. was described by Linnaeus in the 1st edition of his Species Plantarum (1753: 149) indicating: “3. Anagallis foliis cordatis amplexicaulibus, caulibus compressis latifolia/ Anagallis hispanica, latifolia, maximum flore. Tournef. Inst. 142/ Cruciata Montana minor, flore caeruleo. Barr. Ic. 584/ Habitat in Hispania Loefl.”

Linnaeus described that plant indicating the colour of the flowers, “*Corolla caerulea, fondo purpurascens*” and that of the stamens, “*Filamenta purpurea. Antheris oblongis, flavis*”.

The reference made by Linnaeus to Tournefort (1719: 142) corresponds, according to Sampaio (1900: 57), to a plant collected by Tournefort in “Ultra San Joan de Foz ad ostium durii” on his travel through Portugal in 1689, as it appears in a manuscript that Tournefort left in the Museu Botanico da Universidade (Coimbra). This name phrase of Tournefort was used by Sampaio (1900: 58) as a synonym of *Anagallis hispanica* Sampaio, which we consider here synonymous of *Anagallis monelli* L. (see below this species).

The icon 584 of the Lam. 157 by Barrelier (1714: 17), indicated by Linnaeus very faithfully, represents the upper part of a branch with large flowers and with all the lanceolate leaves arranged in whorls of 4 in each node, each with its flower. This plant can also be identified as *Anagallis monelli* L. Therefore, none of the synonyms that Linnaeus gives in *Anagallis latifolia* could be selected as type since both the Tournefort plant and the Barrelier icon have characters very different from those described by Linnaeus in *Anagallis latifolia*. Therefore, the type should be choosen among the material of herbarium that Loefling sent to his Master from Spain, as Linnaeus indicated in the locotypical indication.

In the different Linnaean herbaria, we did not find any material of *Anagallis latifolia* collected by Loefling in Spain. In the main herbarium of Linnaeus (LINN) there is a sheet No. 208.3 with the indication, handwritten by Linnaeus “HU” at the base of the plant and at the base of the sheet “3 latifolia”. 3 is the order that Linnaeus gave to his description of *Anagallis latifolia*, that is, we are dealing with a material that comes from the crops of the Botanical Garden of Uppsala and that was marked by Linnaeus as material of the first edition of his Species Plantarum of 1753, and therefore the material could be the type of the species. In addition, in an attached cut-out to the sheet and handwritten by Linnaeus “Ex Hispanica foly amplexicauly” is indicated. With this statement Linnaeus related the cultivated plant in Hortus Upsaliensis with one from Spain that Loefling sent him. The material that is in the sheet consists of the upper half of a flowered plant of 20 cm, with quadrangular stems and internodes 2-5 cm; opposite leaves, being the bigger 3 x 2 cm, broadly ovate-lanceolate, sharp, cordate at the base; solitary, axillary flowers, with capillary pedicels of 20-30 mm, almost the length of the leaves; blue corolla, with petals of c. 7 mm in length, and immature capsules smaller than the calyx. It is evident that this material comes from cultivation as Linnaeus indicated. Linnaeus also made a detailed description of the species on the living plant, since some characters, especially those related to the colours of flowers and filament and anther of the stamens, cannot be described with material found in ‘Linnaeus’; main herbal sheet. This material from the Uppsala Garden came from seeds that Loefling periodically sent to Linnaeus (López González, 1990: 43). For all the above, we propose as lectotype of *Anagallis latifolia* L. the only plant found in Linnaeus’; main herbarium (LINN) n° 308.3.

-Plants with very small blue flowers that live in wetland environment have been included sometimes in *Anagallis arvensis* thus, a typification proposal is made.

*Lysimachia talaverae* L. Sáez & Aymerich in Orsis 29: 48 (2015) ≡ *Anagallis parviflora* Hoffmanns. & Link, Fl. Portug. 1 (11): 325, 326. Tab. 64 (1813-1820) [syn. subst.] ≡ *Anagallis arvensis* subsp. *parviflora* (Hoffmanns. & Link) Arcangeli, Comp. Fl. Ital. ed.2: 456 (1894) = *Anagallis latifolia* “raza” *parviflora* (Hoffmanns. & Link) Merino in Broteria, sér. Bot. 14: 162 (1916), nom. illeg.

Ind. loc.: “Dans les lieux sablonneux aux environs de Comporta”.

Lectotype designated here (iconotype): *Anagallis parviflora* Tab. 64 in Hoffmannsegg & Link, Fl. Portug. 1 (13): 326 (1813-1820).

Epitype: Huelva. Hinojos, Las Porqueras. Lagoon edge. 37° 17’; 37” N-6° 25’; 15”W. 108 m. 5/5/2014. Leg. F.J. Jiménez & S. Talavera. SEV286467. A sheet of this sample has been used in this study by establish the phylogeny with a nuclear marker (ITS) and three plastic markers.

-*Lysimachia monelli* should be referred exclusively to the blue lineage because it was used by Linnaeus in 1753 to describe this species for the first time.

*Lysimachia monelli* (L.) U. Manns & Anderb. in Willdenowia, 39(1): 52 (2009) ≡ *Anagallis monelli* L., Sp. Pl. 1: 148 (1753), [basion.].

Ind. Loc.: not indicated by Linnaeus in 1753; “Habitat in Verona” in Linnaeus, Sp. Pl. ed. 2, 1: 212. 1762; As indicated Carlos Pau (1915) the locotypical indication was clearly exposed by Linnaeus when he described the species when indicating his previous synonym, both in his Hortus cliffortianus, 52. 1738: “Crecit forte Gadibus und seminis accepit Joh. Monellus Tornacensis, atque eaden cum Clussius comunicavit anno 1602”, as in Hortus upsaliensis, 38. 1748: “Habitario incerta. Johannes Monellus 1662 Gadibus semina Clussius misit. Hospitatur in Terpidario, perennes”.

Lectotype designated by Manns & Anderberg in Willdenowia 39 (1): 52 (2009): Hort. Cliff. 52.2, Anagallis 2 (BM000557969).

The lectotype (photo seen) is composed by two flowering stems of c. 30 cm, one undivided and the other branched almost from the base, with numerous flowers but only those of the apex in anthesis, the others in postanthesis; internodes 2-5 cm, longer than the leaves; opposite inferior leaves, of 20-25 x 5 mm, elliptical, attenuate at the base, acute in the apex, superior in whorls of 3 leaves of 15-20 x 3-5 mm, elliptical, each one with a long pedicelled flower; pedicels of 25-35 mm, generally larger than the leaves. These branches come from the garden of George Clifford III (Hartekamp Garden, Holland).

= *Anagallis linifolia* L. Sp. Pl. ed. 2, 1: 212 (1762)

≡ *Anagallis monelli* subsp. *linifolia* Maire in Jahandiez & Maire, Cat. Pl. Maroc.: 562 (1934)

Ind. Loc.: “Habitat in Lusitania, Hispania. Claud Alstroemer”

Lectotype designated here: “linifolia [m. Linnaeus] A [Alstroemer]”: Herb. Linn. N° 208.4 (LINN-ML)

The lectotype is formed by a single, apparently, annual plant of 10 cm high, erect, with 3 short branches with fruits in the upper half, and other branches emerging near to the axonomorph root; stem with very short internodes, surpassed by the leaves; leaves of 6-10 x 0.1-0.2 mm, narrowly linear, truncated at the base, obtuse; pedicels all with fruits, recurved, rigids; fruits shorter than the calyx. The characters of this plant coincide with those indicated by Linnaeus when he described the species, so there is no doubt that this plant is the same as Linnaeus saw. The indication “Lusitania” could have been made by Linnaeus from a plant from Portugal described by Tournefort and cited as a synonym.

Annual plants such as the type are very frequent on the coast of the province of Cádiz near the capital, so it is not surprising that Linnaeus thought that *A. linifolia* was annual. According to Fernández Pérez (1990), Class Alstroemer toured these territories in the spring-summer of 1760. Probably he collected this plant in late spring when it did not flowers. That is why Linnaeus only described the fruits, as seen in the type material. So the type probably comes from this region of Cádiz.

-Red plants of *Lysimachia monelli* should be called *Lysimachia collina* because plants with this colour were described by Schousboe in 1800 as *Anagallis collina*. Due to *Anagallis* now are included in *Lysimachia*, we do a new combination.

### Lysimachia collina (Schousb.) FJ. Jiménez-López, com.nov

*Anagallis collina* Schousb., Iagttag. Vextrig. Marokko.: 78. (1800), basion. ≡ *Anagallis monelli* subsp. *collina* (Schousb.) H. Lindb. in Acta Soc. Sci. Fenn., ser. B, Opera Biol. 1(2): 115. (1932). ≡ *Anagallis monelli* var. *collina* (Schousb.) Pau in Mem. Real. Soc. Esp.

Hist. Nat. 12 (5): 360 (1924) ≡ *Anagallis linifolia* var. collina (Schousb.) Ball in J. Linn. Soc. 16: 562 (1878). ≡ *Anagallis linifolia* f. rubriflora Batt. in Battandier & Trabut, Fl. Algérie, Dicot.: 723 (1890), [syn. subst.].

Ind. loc.: “Frequens ocurrit in collibus aridis provinciae Haha”

Lectotype designated here: Mogador [Haha] Schousboe [m. Schousboe]: C 10001180.

The type material is composed by a single woody stem of 15 cm length, without leaves in the lower half, branched in the upper half; each branch short, with some oval-lanceolate leaves, acute, shorter than the pedicels and the majority of the flowers in anthesis; isolectotype C 10001181.

## Acknowledgements

We thank the following people for their contribution in the sample collection and data analysis: F. Balao, R. Berjano, M. Buide, I. Casimiro-Soriguer, J.A. Mejías, M.A. Ortiz and S. Talavera. We thank to S. Talavera for comments on the manuscript. This work was funded by Spanish with FEDER funds from the Ministerio de Ciencia e Innovación and MINECO (CGL2008-02531-E, CGL2012-33270 & CGL2015-63827 to M. Arista), the fellowship BES-2013-062859 from MINECO to F.J. Jiménez-López and the VPPI-US to M. Talavera. We thank the Herbarium of the University of Sevilla (CITIUS) for their collaboration and the use of its new DNA Bank, and to Royal Botanic Gardens, Kew.

## Notes

### Competing Interest Statement

The authors have declared no competing interest.

