## Supplemental Table S1 for "Molecular approaches reveal speciation between red and blue flowered plants in the Mediterranean *Lysimachia arvensis* and *Lysimachia monelli* (Primulaceae)"

Table S1. Population identity. For each population is shown country, region, coordinates, flower colour, herbarium code and accession numbers of Genbank database. ID: species and population code. Colour: (B) Blue, (R) Red , (Y) Yellow, (W) White.

| ID | Colour | | Locality | Coordinates | Voucher No. | Genbank Acc. No. | | | |
|---|---|---|---|---|---|---|---|---|---|
| | | | | | | ITS | trnH-psbA | rps16-trnK | rpl32-trnL |
| ***Lysimachia arvensis*** (L.) U. Manns & Anderb. | | | | | | | | | |
| **LA1** | R | PRT | Azores. Sao Jorge | 38°40'35.6"N-28°06'40.8"W | SEV275603 | xxx | xxx | xxx | xxx |
| **LA2** | B | PRT | Albufeira. Praia de Falesia | 37°5'12"N-8º10'9"W | SEV250700 | xxx | xxx | xxx | xxx |
| **LA3** | R | PRT | Madeira. Boaventura | 32º48'55''N-16º58'05''W | SEV285615 | xxx | | | |
| **LA4** | B | ESP | Huelva. Hinojos | 37º17'42.6"N-6º25'26.2"W | SEV279168 | xxx | xxx | xxx | xxx |
| **LA5** | B | ESP | Huelva. Aracena | 37º54'17.6"N-6º34'3.8"W | SEV278862 | xxx | xxx | xxx | xxx |
| **LA5** | R | ESP | Huelva. Aracena | 37º54'17.6"N-6º34'3.8"W | SEV278867 | | xxx | xxx | xxx |
| **LA6** | B | ESP | Huelva. Cañaveral de León | 38°00'50.1"N-6°31'22.9"W | SEV279143 | xxx | | | |
| **LA7** | B | ESP | Badajoz. Monesterio | 38º5'26.9''N-6º17'9.6'' W | SEV279171 | xxx | xxx | xxx | xxx |
| **LA8** | B | ESP | Sevilla. Aznalcázar | 37°18'09.9"N-6°15'45.4"W | SEV250703 | xxx | xxx | xxx | xxx |
| **LA9** | B | ESP | Cádiz. Zahara de los Atunes | 36°06'23.9"N-5°49'34.3"W | SEV279208 | xxx | | | |
| **LA10** | R | ESP | Navarra. Leitza | 43°05'1.04"N-1°54'59.16"W | SEV279272 | xxx | | | |
| **LA11** | B | ESP | Sevilla. El Pedroso | 37°50'16.3"N-5°45'41.9"W | SEV279228 | | | | |
| **LA12** | B | ESP | Cáceres. Losar de la Vera | 40°06'38.0"N-5°34'56.2"W | SEV278765 | | xxx | xxx | xxx |
| **LA13** | B | ESP | Cádiz. Grazalema | 36°45'25.1"N-5°23'42.4"W | SEV279114 | | xxx | xxx | xxx |
| **LA14** | B | ESP | Ávila. Poyales del Hoyo | 40°10'32.6"N-5°09'30.7"W | SEV278770 | | xxx | xxx | xxx |
| **LA14** | R | ESP | Ávila. Poyales del Hoyo | 40°10'32.6"N-5°09'30.7"W | SEV278773 | | xxx | xxx | xxx |
| **LA15** | B | ESP | Málaga. Estepona | 36°25'44.81"N-5°8'4.76"W | - | xxx | xxx | xxx | xxx |
| **LA15** | R | ESP | Málaga. Estepona | 36°25'44.81"N-5°8'4.76"W | SEV285214 | | xxx | xxx | xxx |
| **LA16** | R | ESP | Málaga. San Pedro de Álcántara | 36º29'32.2"N-5º2'27.3"W | SEV286473 | xxx | | | |

| | | | | | | | | | |
|---|---|---|---|---|---|---|---|---|---|
| **LA17** | R | ESP | Córdoba. Carcabuey. Fuente Dura | 37°27'03.8"N-4°16'44.3"W | SEV279258 | xxx | xxx | xxx | xxx |
| **LA18** | B | ESP | Córdoba. Carcabuey | 37º26'23.3"N-4º16'37"W | SEV279276 | xxx | | | |
| **LA19** | B | ESP | Granada. Sierra de Huétor | 37º15'17''N-3º29'8''W | SEV279149 | | | | |
| **LA20** | B | ESP | Mallorca. Parc Natural de Levant | 39º44'10.5"N-3º20'5.5"E | SEV279240 | xxx | xxx | xxx | xxx |
| **LA20** | R | ESP | Mallorca. Parc Natural de Levant | 39º44'10.5"N-3º20'5.5"E | SEV279236 | xxx | xxx | xxx | xxx |
| **LA21** | B | ESP | Alicante. Denia | 38º49'2.6" N-0º6'26.7" E | SEV278776 | xxx | | | |
| **LA22** | R | ESP | Navarra. Oieregi | 43°08'18.3"N-1°37'12.4"W | SEV279266 | xxx | xxx | xxx | xxx |
| **LA23** | B | ESP | Formentera. Es Ca Mari | | SEV252540 | xxx | | | |
| **LA24** | R | ESP | Mallorca. Boal des Ses Severes | 39º38'45.9"N-2º27'49.7"E | SEV256604-1 | xxx | | | |
| **LA25** | R | ITA | Sardinia. Chia | 38º53'38.8"N- 8º51'3.1"E | SEV252545 | xxx | xxx | xxx | xxx |
| **LA26** | B | ITA | Sicily. Scillato-Caltavuturo | 37º50'34.4''N-13º54'14.3''E | SEV279201 | xxx | | | |
| **LA27** | R | TUN | Tabarka. Close to Algerian Frontier | 36º57'46.4''N-8º44'51''E | SEV285272 | | xxx | xxx | xxx |
| **LA28** | B | GRC | Trapeza. Kalávrita | 38°02'08.3"N-22°06'50.5"E | - | xxx | xxx | xxx | xxx |
| **LA28** | R | GRC | Trapeza. Kalávrita | 38°02'08.3"N-22°06'50.5"E | - | xxx | xxx | xxx | xxx |
| **LA29** | B | GRC | Crete. Aglhia Pelaghia | 35º24'36''N-24º59'51''E | SEV279241 | xxx | xxx | xxx | xxx |
| **LA29** | R | GRC | Crete. Aglhia Pelaghia | 35º24'36''N-24º59'51''E | SEV279244 | xxx | xxx | xxx | xxx |
| **LA30** | B | TUR | Antalya. Belek | 36º50'54.5''N-31º4'39.2''E | SEV252544-2 | xxx | xxx | xxx | xxx |
| **LA30** | R | TUR | Antalya. Belek | 36º50'54.5''N-31º4'39.2''E | SEV252544-1 | xxx | xxx | xxx | xxx |
| ***Lysimachia monelli*** (L.) U. Manns & Anderb. | | | | | | | | | |
| **LM1** | B | PRT | Estremadura. Cabo Espichel | 38º25'8''N-9º12'58''W | SEV284843 | xxx | xxx | xxx | xxx |
| **LM2** | B | PRT | Algarve. Sagres | 37º0'20''N-8º56'46''W | SEV284397 | xxx | | | |
| **LM3** | B | PRT | Alentejo Litoral. Sines | 38º0'55.2''N-8º49'11.1''W | SEV284791 | xxx | | | |

| | | | | | | | | | |
|---|---|---|---|---|---|---|---|---|---|
| **LM4** | B | ESP | Huelva. Mazagón | 37º7'26.3''N-6º45'46.2''W | SEV285034 | xxx | xxx | xxx | xxx |
| **LM5** | B | ESP | Cádiz. Zahora | 36º12'11.1''N-6º3'4.9''W | SEV286470 | xxx | | | |
| **LM6** | B | ESP | Cádiz. Barbate | 36º12'21.1''N-5º56'14.3''W | SEV286469 | xxx | | | |
| **LM7** | B | ESP | Salamanca.Béjar | 40º23'47''N-5º49'29''W | SEV285022 | xxx | | | |
| **LM8** | B | ESP | Zamora. Granja de Moreruela | 41º49'49''N-5º43'53''W | SEV284982 | xxx | xxx | xxx | xxx |
| **LM9** | R | ESP | Tarragona. Mont Roig del Camp | 41º5'18''N-0º56'5''E | SEV285107 | xxx | | | |
| **LM10** | R | MAR | El Hajeb-Azrou | 33°31'40.0"N-5°18'25"W | SEV218069 | xxx | xxx | xxx | xxx |
| **LM11** | R | MAR | Fès. J. Bou- Iblane | 33º39'37.1''N-4º14'5.1''W | SEV227757 | xxx | xxx | xxx | xxx |
| **LM12** | R | ITA | Sardinia. Cala Fico | 39º9'17.2''N-8º13'42.5''E | SEV252550 | xxx | xxx | xxx | xxx |
| ***Lysimachia foemina*** **(Mill.) U. Manns & Anderb.** | | | | | | | | | |
| **LF1** | B | ESP | Almeria. Seron | 37°20'55.8"N-2°30'35.3"W | SEV269286 | xxx | xxx | xxx | xxx |
| **LF2** | B | ESP | Málaga. Almargen-Cañete la Real | 36°58'6.56"N-5°2'33.18"W | SEV285199 | xxx | | | |
| **LF3** | B | ESP | Mallorca. Boal des Ses Severes | 39º38'45.9"N-2º27'49.7"E | SEV256604-2 | xxx | | | |
| ***Lysimachia*** **talaverae** L. Sáez & Aymerich | | | | | | | | | |
| **LT1** | B | ESP | Huelva. Almonte | 37°08'39.9"N-6°32'57.6"W | SEV286468 | xxx | xxx | xxx | xxx |
| **LT2** | B | ESP | Huelva. Hinojos | 37º17'37''N-6º25'15''W | SEV286467 | xxx | xxx | xxx | xxx |
| **LT3** | B | PRT | Algarve. Vila do Bispo | 37º7'18''N-8º53'39''W | SEV284451 | xxx | | | |
| **LT4** | B | PRT | Ribatejo. Coruche | 38º50'8''N-8º33'31''W | SEV284877 | xxx | | | |
| ***Lysimachia azorica*** **Hornem. ex Hook.** | | | | | | | | | |
| **LZ1** | Y | PRT | Azores. Terceira. Algar do Carvao | 38º43'54.4''N-27º19'16.7''W | SEV275597 | xxx | xxx | xxx | xxx |
| ***Lysimachia linum-stellatum*** **(L.) Duby** | | | | | | | | | |
| **LLS1** | W | ESP | Sevilla. Dos Hermanas | 37°21'06.5"N-5°56'21.9"W | SEV286828 | xxx | | | |
| **LLS1a** | W | ESP | Sevilla. Dos Hermanas | 37°21'06.5"N-5°56'21.9"W | SEV286828 | xxx | | | |

| | |
|---|---|
| ***Lysimachia tyrrhenia*** U. Manns & Anderb. (former *Anagallis crassifolia* Thore) | |
| | GenBank Acc. No: AY855136 |
| ***Lysimachia tenella*** L. (former *Anagallis tenella* L.) | |
| | GenBank Acc. No: AY855150 |
